# Identification of candidate genes for gelatinization temperature, gel consistency and pericarp color by GWAS in rice based on SLAF-sequencing

**DOI:** 10.1101/258244

**Authors:** Xinghai Yang, Xiuzhong Xia, Yu Zeng, Baoxuan Nong, Zongqiong Zhang, Yanyan Wu, Faqian Xiong, Yuexiong Zhang, Haifu Liang, Guofu Deng, Danting Li

## Abstract

Rice is an important cereal in the world, uncovering the genetic basis of agronomic traits in rice landraces genes associated with agronomically important traits is indispensable for both understanding the genetic basis of phenotypic variation and efficient crop improvement. Gelatinization temperature, gel consistency and pericarp color are important indices of rice cooking and eating quality evaluation and potential nutritional importance, which attract wide attentions in the application of genetic and breeding. To dissect the genetic basis of gelatinization temperature (GT), gel consistency (GC) and pericarp color (PC), a total of 419 rice landraces core germplasm collections consisting of 330 *indica* lines, 78 *japonica* lines and 11 uncertain varieties were grown, collected, then GT, GC, PC were measured for two years, and sequenced using Specific Locus Amplified Fragment Sequencing (SLAF) technology. In this study, 261,385,070 clean reads and 56,768 polymorphic SLAF tags were obtained, which a total of 211,818 single nucleotide polymorphisms (SNPs) were discovered. With 208,993 SNPs meeting the criterion of minor allele frequency (MAF) > 0.05 and integrity> 0.5, the phylogenetic tree and population structure analysis were performed for all 419 rice landraces, and the whole panel mainly separated into six subpopulations based on population structure analysis. Genome-wide association study (GWAS) was carried out for the whole panel, *indica* subpanel and *japonica* subpanel with subset SNPs respectively. One quantitative trait locus (QTL) on chromosome 6 for GT was detected in the whole panel and *indica* subpanel, and one QTL associated with GC was located on chromosome 6 in the whole panel and *indica* subpanel. For the PC trait, 8 QTLs were detected in the whole panel on chromosome 1, 3, 4, 7, 8, 10 and 11, and 7 QTLs in the *indica* subpanel on chromosome 3, 4, 7, 8, 10 and 11. The loci on chromosome 3, 8, 10 and 11 have not been identified previously, and they may be the candidate genes of pericarp color. For the three traits, no QTL was detected in *japonica* subpanel probably because of the polymorphism repartition between the subpanel, or small population size of *japonica* subpanel. This paper provides new gene resources and insights into the molecular mechanisms of important agricultural trait of rice phenotypic variation and genetic improvement of rice quality variety breeding.

## Introduction

Rice is one of the most important food crops in the world[1], the increased rice yield, improved quality and advanced resistance to biotic and abiotic stress play an important role in solving the world food problem, improving people’s life quality and reducing environmental pollution. Identification and utilization of favorable genes in rice germplasm resources is the foundation of rice breeding. Guangxi is likely the origin of cultivated rice[2], which possesses a large number of rice landraces containing rich natural variation and genetic diversity, and it is the important genetic resources for breed improvement.

Quantitative trait loci (QTLs) mapping has been widely used to explore the genetic basis of complex agronomic traits in different crops. So far, yield-related genes [3–6], quality-related genes [7–9], and resistance-related genes [10–12] and so on have been identified and cloned through biparental linkage mapping in rice (http://www.gramene.org/, http://www.ricedata.cn/index.htm). However, almost studies need to construct mapping populations (e.g. F2, BC, RILs, NILs), which is very time-consuming and painstaking [13].

In recent years, with the rapid development of high-throughput sequencing technology and reduced sequencing cost, genome-wide association study (GWAS) based on SNPs has become a new method for studying important agronomic traits in rice [14–20], maize [21], sorghum [22], soybean [23], tomato [24], et al. Previous studies have documented that GWAS could be a useful tool to dissect the genetic changes for complicated traits has some advantages. First, GWAS can efficiently detect multiple QTLs in the same population meanwhile; Second, the associated population includes the majority of variations of the related loci. Therefore, the GWAS not only can help us understand the gene function, but also discover the favorable alleles for genetic improvement of plants. However, GWAS has some limitations. (i) the large effect variations and minor effect genes can not be identified easily by GWAS [25]; (ii) genetic heterogeneity can reduce the efficacy of mutation detection[25,26]; (iii) the population structure lead to false positive associations between phenotype and unlinked markers [18]; (iv) GWAS can be limited by the genetic characteristics of different species, for example, the LD decay of rice is lower than that of outcrossing maize [27], so the GWAS still can not replace the traditional map based clone to achieve fine mapping of the target gene [18,28].

Although a number of GWAS analysis for important agronomic traits in rice have been performed, new QTLs and candidate genes can be found through different populations [14–20]. In this study, we used 419 rice landraces core germplasm collections from Guangxi to explore the molecular basis of GT, GC and PC, the panel was genotyped based on SLAF-seq technology, the three traits were measured for twice during 2014 and 2015, and then the GWAS was performed. The research provided genetic resources for molecular breeding and shed light on rice quality improvement.

## Materials and methods

### Plant materials and phenotyping

The diversity panel was composed of 419 landraces collected from core rice germplasm of Guangxi. The lines were planted in Nanning experimental field (China at 22.85^o^N, 108.26^o^E) from July 2014 to November 2014 and from July 2015 to November 2015. For each landrace, five randomly chosen plants were harvested when they were mature and used for measurement of gelatinization temperature and gel consistency and recording the pericarp color. The gelatinization temperature was measured by the alkali digestion test [29] and the gel consistency was estimated based on Cagampang et al. (1973) [30].

### SLAF sequencing and SNP genotyping

Total genomic DNA was extracted from young quadrifoliate leaves of all rice landraces using cetyltrimethyl annonium bromide (CTAB) protocol [31], and digested by two restriction enzymes RsaI and HaeIII. The SLAF sequencing was performed on an IlluminaHiseq 2500 system. The polymorphic SLAF tags were obtained by clustering the clean reads using BLAT software [32], aligned to reference genome (*Oryza sativa* L. spp. *japonica*. cv. Nipponbare, http://plants.ensembl.org/index.html) using the BWA software [33], and then the SNP calling was performed using GATK [34] and Samtools packages [35]. A total of 208,993 SNPs with a minor allele frequency (MAF)>0.05 and integrity > 0.5 was retained for GWAS.

### Genetic kinship calculation, phylogenetic tree construction, principal component analysis, population sturcture

Based on the 208,993 high quality SNPs, the phylogenetic tree was constructed by MEGA5 [36]. the pairwise kinship was carried out using the SPAGeDi software package [37], and the population structure was analysed by Admixture software [38], which the subpanel number was predicted from 1 to 10. The kinship and population structure analysis was performed for the whole panel, *indica* subpanel and *japonica* subpanel respectively in convenience of subsequent GWAS for the three combinations. Principal component analysis (PCA) was performed using GAPIT [39].

### Genome-wide association study

In this study, in order to eliminate the effort of population structure between *indica* subpopulation and *japonica* subpopulation, GWAS was proceeded for the whole panel, *indica* subpanel and *japonica* subpanel respectively using the mixed linear model (MLM) of Tassel v3 [40], which took the population structure and kinship into consideration, and the significant P value was set to 4.79×10^-8^.

## Result

### Phenotypic variation of gelatinization temperature, gel consistency and pericarp color

The alkali spreading value can be used to measure the gelatinization temperature, which is inversely related to GT, and ranged from 0 to 6.15 with an average of 2.86 in 2014, and 0 to 7 with an average of 3.74 in 2015; The gel consistency spanned 26 to 100 with an average of 67.91 in 2014, and 6 to 100 with an average of 70.94 in 2015; for the pericarp color trait, the number of rice landraces with white, red, black color is 308, 97, 14 respectively ( S1 Table).

### Analysis of SLAF-seq data and development of SNPs

After sequencing data quality control, a total of 67,665 SLAF tags were obtained with an average sequencing depth of 8.75×. Ultimately, 56,768 polymorphic SLAF tags were retained when aligned to the reference genome (S2 Table).

A total of 211,818 SNPs was identified using the GATK and samtools software package, of these, the number of SNPs with MAF > 0.05 and integrity> 0.5 was 208,993, which distributed on every chromosome with a mean number 17,416 per chromosome. For the high quality SNPs, 104,068 SNPs located in gene region, and 107750 SNPs in intergenic region. The chromosome 10 possessed the most SNP density while chromosome 2 for the fewest SNP density (Fig 1, S3 Table).

### Genetic kinship calculation, phylogenetic tree construction and principal component analysis

Based on the 208,993 high quality SNPs, 87,571 pairwise calculations for all 419 rice landraces were carried out. Of these, the number of pairs reached up to 45,709, which the genetic relationship coefficients <0.05. The phylogenetic tree was clustered two mainly panels in accordance with the *indica* and *japonica* subpopulations. The first two principal components explained 5.64% and 3.92% of the genetic variation respectively (Fig 2).

### Analysis of population sturcture

Based on the error rate of 5-fold cross-validation, the ancestor number was confirmed to 6 for all the 419 rice landraces. The six subpanels respectively contain 330 *indica* varieties, 78 *japonica* varities (Fig 3).

### GWAS mapping in rice landraces

The whole panel composed of 419 rice landraces, *indica* subpanel (330 rice landraces) and *japonica* subpanel (78 rice landraces) were utilized for GWAS respectively in order to avoid population structure noise. Only one QTL was detected for GT, but in both the whole population and *indica* subpopulation. Similarly, only one association with GC was obtained, also both the whole population and *indica* subpopulation. Interesting, the two QTLs were both located on chromosome 6 (Fig 4, Fig 5). For the pericarp color trait, 8 QTLs were detected in the whole panel on chromosome1, 3,4,7, 8, 10 and 11, and 7 QTLs in the *indica* subpanel on chromosome 3, 4, 7, 8, 10 and 11 (Fig 6).

### GWAS on Gelatinization Temperature

Gelatinization temperature is one of the most important indexes to evaluate the cooking and eating quality of rice. The association analysis for GT was conducted for the whole panel, *indica* subpanel and *japonica* subpanel successively. In 2014, for the whole panel of 419 rice accessions, GWAS detected a total of 48 GT related SNPs, the 26 SNPs of which can also be detected in *indica* subpanel, and the significant associated SNPs distributed on the 1807797 bp-7174281 bp of chromosome 6 (Fig 4, S4 Table). In the whole and *indica* panels, the most significant associated SNPs with GT were Chr6_6733351 (P=2.04x10E-15) and Chr6_6740370 (1.39x10E-13). In 2015, 8 GT related SNPs were identified in the whole panel, of which 4 SNPs can also be detected in *indica* panel, and the positions of SNPs ranged from 6740370 bp to 6927719 bp of chromosome 6 (Fig 4, S4 Table). However, no significant SNPs were detected for the *japonica* subpanel in the both 2014 and 2015 year. In the whole panel and *indica* subpanel, the most significant associated SNPs were both Chr6_6879531 (2.23x10E-11, 1.14><10E-9). Chr6_6733351 and Chr6_6740370 were located in 15kb and 8 kbupstream of the *ALK*gene (*LOC_Os06g12450*), respectively (Table 1).

**Table 1.**
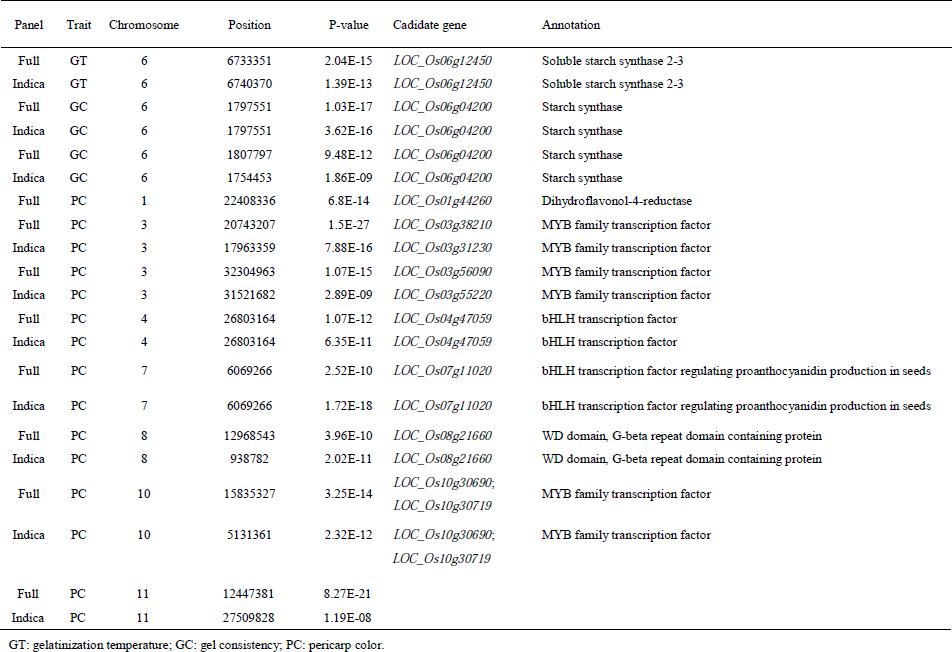
A sbuset of associated loci and candidate genes.

The *ALK* gene encodes soluble starch synthase a *SSIIa* and is responsible for GT of rice [7]. Some researchers have analyzed the SNP in *ALK* gene, and considered that SNP variation is the main factor of gelatinization temperature change [41–43]. We also found a SNP Chr6_1807797 (P=2.53×10^-8^) which was significantly associated with GT located in the *Wx* gene region in 2014, which further verified the correlation between the *ALK* gene and the *Wx* gene (Fig 4).

### GWAS on Gel Consistency

Gel consistency is a complex quality trait, and views about the genetic basis of GC were not consistent between researchers (http://www.gramene.org/). Gel consistency is inversely related to amylose content, we performed Pearson correlation analysis for gel consistency and amylose content trait, and reached the same conclusion (S5 Table). Many researchers believed that the *Wx* gene located on the Chromosome 6 of rice is the major gene controlling the gel consistency [44–46], and some other GC-related QTLs located on chromosome 1, 2, 3, 6, and 7 were also detected [47,48]. In 2014, in the whole panel and *indica* subpanel, GWAS detected a total of 28 and 24 significantly associated SNPs respectively, the QTL located in the interval of 1607061 bp-1958767 bp of chromosome 6, and the most significant associated SNP were both Chr6_1797551 (P=1.03x10E-17; P=3.62x10E-16) for both whole and *indica* panels (Fig 5, S6 Table). In 2015, 13 and 4 associated SNPs were confirmed for the whole and *indica* panels respectively. The associated SNPs sited in 1661801 bp-1822395 bp of chromosome 6 (Fig 5, S6 Table). However, no SNPs were detected for the *japonica* subpanel in both 2014 and 2015 year. For the whole and *indica* panels, the most significant associated SNPs were Chr6_1807797 (P=9.48x10E-12) and Chr6_1754453 (P=1.86x10E-9) respectively. The Chr6_1754453, Chr6_1797551 and Chr6_1807797 were located 11.2kb upstream, 26.9kb downstream and 37.2kb downstream of the *Wx*gene (*LOC_Os06g04200*) respectively (Table 1).

### GWAS on Pericarp Color

Proanthocyanidins and anthocyanins are accumulated in the red pericarp rice and black pericarp rice respectively. The anthocyanin biosynthesis pathway includes multiple structural genes, such as *CHS, CHI* and *DFR*. A large number of studies have shown that these structural gene expressions have different degrees of synergy, which are directly controlled by MBW protein complexes formed by MYB, bHLH and WD40 transcription factors. The synthesis of anthocyanin in most plants is regulated by MBW through binding to the promoter of structural genes.

For the whole and *indica* panels, 763 and 99 significantly associated SNPs were identified respectively, however significant SNPs were not detected for *japonica* panel (Fig 6, S7 Table). For chromosome 1 of rice, 25 significantly associated SNPs were detected in the whole panel, but no SNPs in *indica* subpanel. The most significant SNP was Chr1_22408336 (P=6.80x10E-14), located in the 2.97 Mb downstream of proanthocyanidins biosynthesis gene *Rd*[49] (*LOC_Os01g44260)*. For chromosome 3, 647 and 27 significant associated SNPs were detected for the whole and *indica* pane respectively, which sited in two clearly divided QTLs. In the 17126203 bp-24432074 bp, the most significantly associated loci were Chr3_20743207 (P=1.50x10E-27) and Chr3_17963359 (P=7.88x10E-16) for the whole and *indica* panels respectively. The *LOC_Os03g38210* and *LOC_Os03g31230*, which both encode the MYB family transcription factors, were located in the 459.6 kb downstream of Chr3_20743207 and 179.2kb upstream of Chr3_17963359 respectively (Table 1). In the 31984778 bp-34408499 bp of chromosome 3, for the whole and *indica* panels, the most significantly associated SNPs were Chr3_32304963 (P=1.07x10E-15) and Chr3_31521682 (P=2.89x10E-9); the Chr3_32304963 sited in the 350.3kb downstream of gene *LOC_Os03g56090* encoding MYB family transcription factor; Chr3_31521682 located in 94.8kb downstream of *LOC_Os03g55220* and 86.9kb downstream of *LOC_Os03g55550*, which both encode bHelix-loop-helix transcription factor (Table 1). For chromosome 4, 11 and 5 significantly associated SNPs were found between whole and *indica* panels, and the most significant SNPs were both Chr4_26803164 (P=1.07x10-E12, P=6.35x10E-12) (Table 1). *Kala4* was located in the 1.11 Mb downstream of Chr4_26803164, which encodes a bHLH transcription factor, provided structural rearrangements of its promoter region, then resulting in its ectopic expression, thus giving the trait of black pericarp in rice [50]. For chromosome 7, 12 significant SNPs were detected in the whole panel, and 61 significant SNPs in *indica* subpanel. The most significant SNP was Chr6_6069266 (2.52x10E-10; 1.72x10E-18), located in the *Rc* gene region (Table 1), which encodes a bHLH motif contained protein that participates in the synthesis of proanthocyanidins of pericarp, and the 14-bp deletion of seventh exon of Rc leads to the Rc mutation to *rc* [51]. For chromosome 8, 17 significantly associated SNPs for the whole panel and 2 SNPs for *indica* subpanel were detected. For the whole panel, the gene *LOC_Os08g21660* encoding a WD domain, G-beta repeat domain contained protein located in 59.9kb upstream of Chr8_12968543 (P=3.96x10E-10) (Table 1), which also located in the QTL ranging from 938782 bp to 20998896 bp of *indica* subpanel. For chromosome 10, 37 significant SNPs for the whole panel and 3 significant SNPs for *indica* panel were found, and there was no identical SNPs between them. Forthe whole panel, *LOC_Os10g30690* and *LOC_Os10g30719* (Table 1), which encode MYB family transcription factor, were located in 148.5kb and 162.3kb downstream of the most significant SNP Chr10_15835327 (P=3.25x10E-14) respectively. For Chr11, 14 and 1 significant SNPs were detected for the whole and *indica* panels respectively, but no candidate genes related to the pericarp color were found.

## Discussion

### SNP markers obtained by SLAF-Seq technology

Rice is the most important staple crop worldwide, and feeding a fast growing population, so it is emergent to identify genes related to agronomically important traits.The association analysis is powerful to identify phenotypic variance related nucleotide polymorphisms [52,53]. In this study, in order to dissect the genetic basis of gelatinization temperature, gel consistency and pericarp color, we performed GWAS based on SLAF-seq technology. A total of 67,665 SLAF tags were obtained with an average sequencing depth of 8.75x. Ultimately, 56,768 polymorphic SLAF tags were retained when aligned to the reference genome, which polymorphic ratio reached up to 83.89%. A total of 211,818 SNPs were identified and the number of SNPs with MAF > 0.05 and integrity > 0.5 was 208,993, which then were used for association analysis. Wang et al. (2016) [68] have documented that low-coverage whole-genome sequencing is an effective strategy for genome-wide association studies in rice. Our research also confirms the conclusion that the SLAF-technology can effectively and accurately identify the associated genes, though the obtained genome information is less than information, which is obtained by whole genome resequencing.

### QTL comparison among different panels

In this study, in order to eliminate the noise of population structure, we conducted association analysis for the whole panel, *indica* subpanel and *japonica* subpanel respectively. The GWAS results in 2014 and 2015showed that the QTLs of GT, GC and PC were relatively stable in different environmental conditions. For gelatinization temperature, one quantitative trait locus on chromosome 6 was detected in the whole panel and *indica* subpanel, which overlapped with *ALK* gene (Table 1), a confirmed major gene for GT [7]. One association with gel consistency was located on chromosome 6 in the whole panel and *indica* subpanel, where *Wx (LOC_Os06g04200*) located (Table 1, http://rice.plantbiology.msu.edu/). For the pericarp color trait, eight QTLs were detected in the whole panel on chromosome 1, 3, 4, 7, 8, 10 and 11, and seven QTLs in the *indica* subpanel on chromosome 3, 4, 7, 8, 10 and 11 (Table 1). For the three traits, no QTL was detected in *japonica* subpanel probably because of the polymorphism repartition between the subpanel [54], or small population size of *japonica* subpanel [25].

### Analysis of candidate genes for GT, GC and PC

Core collection is a subset of the germplasm resources, representative of the genetic diversity and geographical distribution of the entire population with the minimum number of genetic resources [55]. Guangxi province of China is likely to be the origin of cultivated rice [2], which possesses a large number of rice germplasms and genetic resources.

In this study, a GWAS for GT, GC and PC of 419 rice landraces core germplasms from guangxi was performed, and it was concluded that *ALK* is the major effect gene for GT [56,57] and *Wx* has a minor effect on gelatinization temperature (2009) [58]. GC is an eating and cooking quality related trait with complicated genetic basis. So far, more than 20 QTLs for gel consistency have been detected (http://www.gramene.org) in rice, which located on chromosome1, 2, 4, 7, 11. Many researchers asserted that *Wx* is the major gene for GC [7,45,46,56]. Swamy et al. (2012) [59] identified 6 QTLs for GC located on chromosome 1, 2, 4, 11 in BC2F2 population. Based on RAD-seqtechnology, Peng et al. (2016) [48] confirmed a *Wx* linkaged QTL for GC on chromosome 6 in BC1F5 population derived from crosses between YVB x V20B. Li et al. (2004) [47] found two QTLs for GC on chromosome 2, 7 through RFLP and SSR marker in BC3F1 population obtained from V20A (O. *sativa* L.) x103544 (O. *glaberrima* S.). In this study, the major effort QTL for GC located on chromosome 6, which overlapped *Wx* gene.

The accumulation of proanthocyanidins and anthocyanins in the testa leads to red and black pericarp respectively. The anthocyanin metabolism pathway in *Zea mays, Antirrhinum majus* and *Arabidopsis thaliana* has been explored to some extent [60], which is not yet fully understood in rice. So far, some structural genes involved in anthocyanin biosynthesis has been identified, such as *OsCHS*1 [61], *OsCHS2* [62], *OsCHI* [63], *OsF3H* [64], *OsF3’H* [62], *OsDFR(Rd*) [49] and *OsANS* [62]. However, the regulatory genes, MYB, bHLH and WD40 are not fully reported. Furukawa et al. cloned *Rc* gene located on chromosome 7, which encodes bHLH domain contained transcription factor and participates in the proanthocyanidin synthesis [51], which is consistent with our result. Oikawa et al. (2015) [50] identified *Kala4* gene controlling anthocyanin biosynthesis, identical to *OSB2* gene confirmed by Sakamoto et al. (2001) [65] in rice. Chin et al.(2016) [66] have showed that the *OsC1* gene of chromosome 6 is related to the purple sheath in rice, encoding a MYB transcription factor. In this study, for the whole panel, a pericarp color-related QTL was detected, where the most significant SNP Chr1_22408336 was adjacent to the *Rd* [49] participating in the proanthocyanidins biosynthesis. The *LOC_Os03g38210*, *LOC_Os03g31230* and *LOC_Os03g56090* all encode a MYB transcription factor, which are different locus of *Kala3 Os03g0410000* participating in anthocyanin biosynthesis for black rice [67]. Both *LOC_Os03g55220* and *LOC_Os03g55550* encode bHLH motif contained transcription factor. The anthocyanin biosynthesis related key gene *Kala4* locates in the significant QTL of chromosome 4 [50]. The *LOC_Os08g21660* gene locates in 54.35kb upstream of Chr8_12968543, and encodes a WD domain and G-beta repeat domain contained protein. So far, the WD40 domain contained gene has not been identified in rice, which is involved in the anthocyanin biosynthesis [50]. There are two *genes(LOC_Os10g30690* and *LOC_Os10g30719*) located in 148.5 kb and 162.3 kb downstream of most significant SNP of Chromosome 10 respectively, both encoding MYB transcription factor, which are not overlapped the gene *qPc10 (Os10g0536400*) identified by Wang et al. (2016) [68].

With the improvement of people’s living standard, the rice of high quality is more and more needed. The discovery and utilization of excellent germplasm can accelerate rice breeding. Based on SLAF-seq technology, GWAS for gelatinization temperature, gel consistency and pericarp color of 419 rice core collections from Guangxi was conducted, and associated genes, especially the anthocyanin synthesis related genes on chromosome 3,8,10 and 11 were reported for the first time. This study shed light on the genetic analysis for important agricultural trait of rice and beneficial to plant breeder.

## Acknowledgments

This study was financially supported by National key R&D projects (2016YFD0100101-03), Guangxi’s Ministry of Science and Technology (AB16380117), Guangxi Natural Science Foundation of China (2015GXNSFAA139054) and Guangxi Academy of Agricultural Sciences (2015YT15; 2016JM09).

## Author Contributions

Xinghai Yang designed, performed the experiment and whrote the manuscript, Xiuzhong Xia, Yu Zeng, Baoxuan Nong, Zongqiong Zhang performed the experiment, Yanyan Wu, Faqian Xiong, Yuexiong Zhang, Haifu Liang, Guofu Deng collected and analyzed data, Li Danting designed and revised the manuscript. All authors reviewed and approved this submission.

